# Discovery of small molecule antagonists of the USP5 zinc finger ubiquitin-binding domain

**DOI:** 10.1101/676668

**Authors:** Mandeep K. Mann, Ivan Franzoni, Renato Ferreira de Freitas, Wolfram Tempel, Scott Houliston, Cheryl H. Arrowsmith, Rachel J. Harding, Matthieu Schapira

## Abstract

USP5 disassembles unanchored polyubiquitin chains to recycle free mono-ubiquitin, and is one of twelve ubiquitin-specific proteases featuring a zinc finger ubiquitin-binding domain (ZnF-UBD). This distinct structural module has been associated with substrate positioning or allosteric modulation of catalytic activity, but its cellular function remains unclear. We screened a chemical library focused on the ZnF-UBD of USP5, crystallized hits in complex with the protein, and generated a preliminary structure-activity relationship which enables the development of more potent and selective compounds. This work serves as a framework for the discovery of a chemical probe to delineate the function of USP5 ZnF-UBD in proteasomal degradation and other ubiquitin signalling pathways in health and disease.

## Introduction

Ubiquitination is a reversible, post-translational modification involving the conjugation of a conserved 76 amino acid protein, ubiquitin (Ub), to substrate proteins for proteasomal targeting or the regulation of cell signaling^1–3^. Substrate proteins are generally ubiquitinated through an isopeptide bond between the C-terminal glycine residue of ubiquitin and the terminal nitrogen of lysine side chains^4–6^. Substrate proteins can be mono-or poly-ubiquitinated, where ubiquitin chains are formed through isopeptide linkages with any one of seven internal lysine surface residues or conjugation to the N-terminal amine group^7^. Deubiquitination, the removal of ubiquitin, is carried out by a family of deubiquitinase enzymes (DUBs)^8–10^. Ubiquitin specific proteases (USPs) are the largest sub-family of DUBs consisting of more than 50 cysteine-proteases with diverse roles in ubiquitin homeostasis^3^.

USP5, also known as isopeptidase T or isoT, disassembles a variety of unanchored polyubiquitin chains (Lys6, Lys11, Lys27, Lys29, Lys33, Lys48, Lys63)^11,12^ to recycle free mono-ubiquitin. USP5 prevents accumulation of poly-ubiquitin chains which would otherwise overwhelm the proteasome by competing with ubiquitinated proteins targeted for degradation^13^. The domain architecture of USP5 consists of a zinc finger ubiquitin-binding domain (ZnF-UBD) spanning residues 173-283, a functional catalytic domain, two ubiquitin associated domains (UBA), and a cryptic zinc finger ubiquitin binding domain (nUBP)^12^. The ZnF-UBD recognizes the C-terminal di-glycine motif of ubiquitin^11^, and has been associated with allosteric modulation of USP5 activity^8,11,14^, and alternatively, with substrate recognition and positioning^12^.

USP5 has been associated with tumorigenesis^15–19^, neurodegeneration^20–22^, and viral infections^23,24^. In addition, USP5 plays a role in regulation of ubiquitin levels in DNA damage repair^25^ and stress granules^26^. According to CRISPR-knockout screens against large panels of cancer cell lines, USP5 is essential to the survival of cells from most cancer types^27–29^. Inhibition of USP catalytic activity by small molecules has been reported for USP1, USP7 and USP14^30–34^. Targeting the non-catalytic ZnF-UBD may be an alternative strategy to antagonize USP5 function, similar to the recent development of small molecule inhibitors against the non-catalytic domain of HDAC6^35,36^. Here, we report the structure-based discovery of the first USP5 ZnF-UBD inhibitors and their characterization by ^19^F NMR spectroscopy, surface plasmon resonance (SPR) and X-ray crystallography. Experimental validation of four low-affinity ligands followed by a hit expansion campaign defined a preliminary structure activity relationship (SAR). Our work provides a framework for the development of potent USP5 ZnF-UBD inhibitors to investigate the function of this non-catalytic domain in cells.

## Results and Discussion

### Virtual Screen & Hit Identification

Following our observation that compounds featuring a short carboxylic acid chain can mimic the C-terminal di-glycine carboxylate of ubiquitin and favorably exploit the ZnF-UBD of HDAC6^35,36^, we compiled a virtual library of 9480 in-house and commercially available compounds to screen against USP5 ZnF-UBD. The library was docked using ICM-Pro^37^ (Molsoft, CA) and Glide^38^ (Schrodinger, NY) to the X-ray crystal structure conformation of USP5 ZnF-UBD (PDB: 2G45) and an alternate conformational state modelled after the structure of HDAC6 ZnF-UBD in complex with inhibitors. In this alternate “stacked” conformation, the guanidinium plane of Arg221 is positioned parallel to the phenyl plane of Tyr259 (Figure S1), which would allow favorable π-stacking interaction with aromatic inhibitors. 96 compounds were selected from ICM^37^ and Glide^38^ by visual inspection of the docked binding pose and physical samples of 33 (mostly fragments) were tested experimentally. Since one of the two tryptophan side-chains of USP5 ZnF-UBD is located at the targeted ubiquitin C-terminal binding pocket (Figure 1a), we decided to use a ^19^F nuclear magnetic resonance (NMR) spectroscopy assay as a primary, qualitative screening method. We first verified that USP5 ZnF-UBD labeled with 5-fluoro-tryptophan (5FW) produced two distinct peaks in a ^19^F spectrum. Titration of 5FW-USP5 ZnF-UBD with the C-terminal ubiquitin peptide LRLRGG resulted in broadening of the downfield peak (Figure 1b), which we consequently assigned as Trp209 in the ubiquitin-binding site. Upon screening the 33 compounds, we found that 11 caused chemical shift perturbation and/or broadening of ^19^F resonance, suggesting binding at the ubiquitin binding pocket (Figure 1c, Table S2). We next used SPR as an orthogonal binding assay to validate primary hits (Figure 1d, Table 1). While the LRLRGG ubiquitin peptide bound USP5 ZnF-UBD with a K_D_ of 52 ± 2 µM, 7 of the 11 NMR hits showed weak affinities in the high micromolar range (Table 1). Fragments **5** and **7** were the most potent ligands with a K_D_ of 220 ± 23 µM, 170 ± 50 µM, and ligand efficiency (LE) of 0.33 and 0.30, respectively (Figure 1d).

**Table 1.**
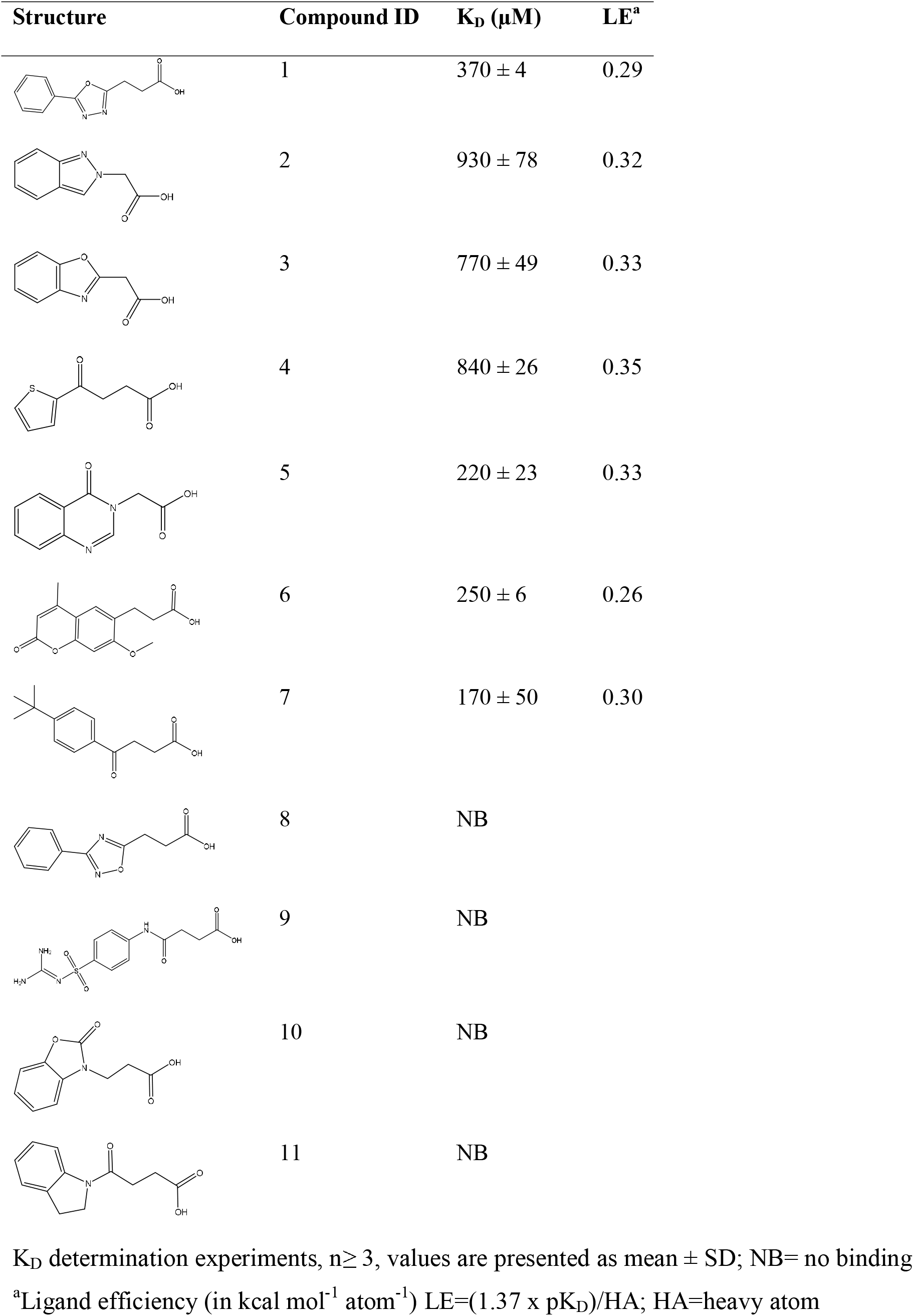
SPR K_D_ and Ligand Efficiency of Preliminary Hit Compounds

| Structure | Compound ID | $K_D$ ($\mu\text{M}$) | LE<sup>a</sup> |
| --- | --- | --- | --- |
| | 1 | $370 \pm 4$ | 0.29 |
| | 2 | $930 \pm 78$ | 0.32 |
| | 3 | $770 \pm 49$ | 0.33 |
| | 4 | $840 \pm 26$ | 0.35 |
| | 5 | $220 \pm 23$ | 0.33 |
| | 6 | $250 \pm 6$ | 0.26 |
| | 7 | $170 \pm 50$ | 0.30 |
|  | 8 | NB |  |
|  | 9 | NB |  |
|  | 10 | NB |  |
|  | 11 | NB |  |

$K_D$ determination experiments, $n \geq 3$ , values are presented as mean ± SD; NB= no binding
<sup>a</sup>Ligand efficiency (in kcal mol<sup>-1</sup> atom<sup>-1</sup>) LE=(1.37 x pK<sub>D</sub>)/HA; HA=heavy atom

**Figure 1.**
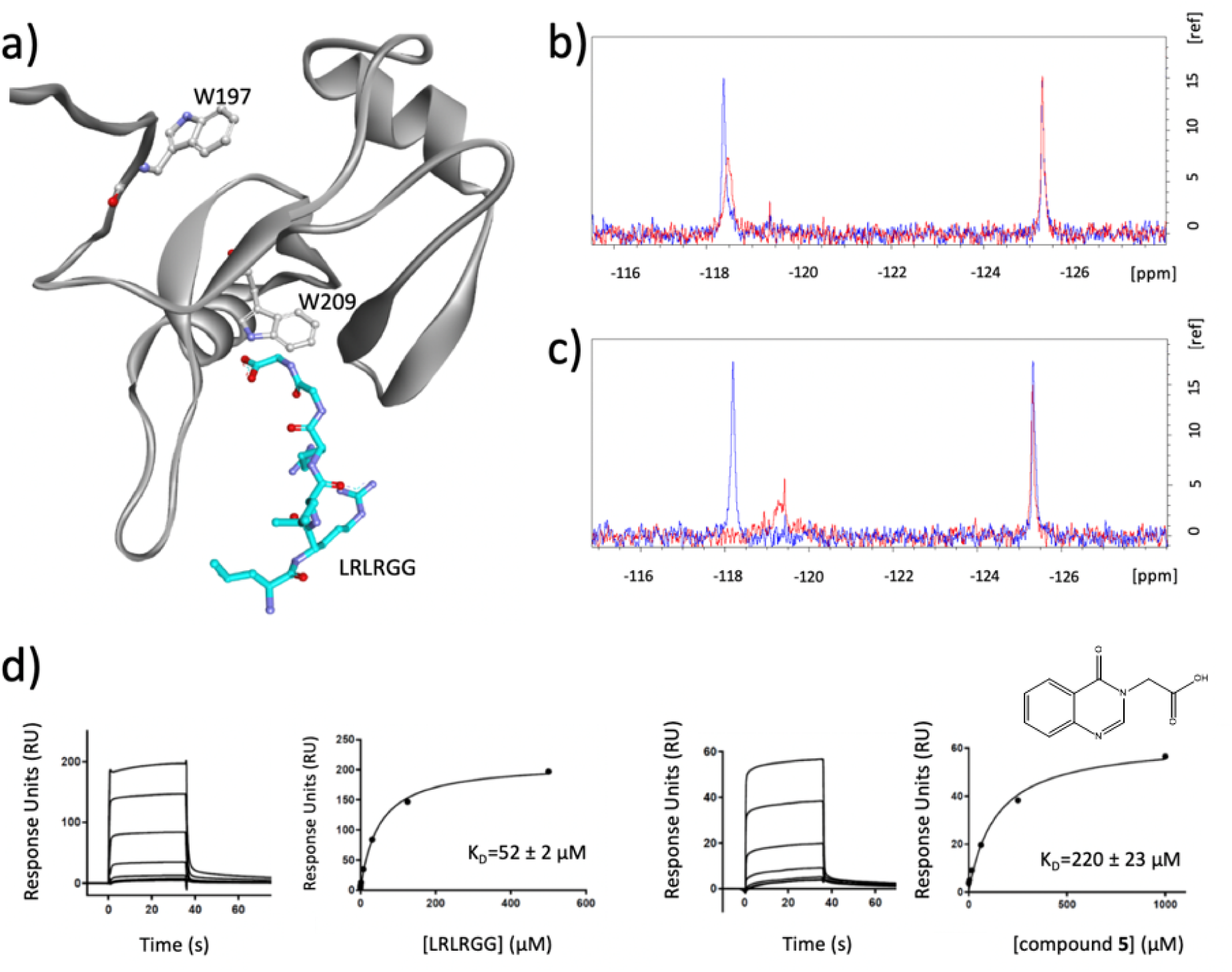
a) Crystal structure of USP5 ZnF-UBD (grey) in complex with a C-terminal ubiquitin peptide (cyan) (PDB: 2G45). The Trp side-chains of USP5 are highlighted. b) ^19^F NMR spectra of 5FW-USP5 ZnF-UBD before (blue) and after the addition of LRLRGG ubiquitin peptide (red) (at 6-fold excess) c) ^19^F NMR spectra of 5FW-USP5 ZnF-UBD before (blue) and after the addition of **5** (at 25-fold excess) (red) d) SPR sensogram and binding curve of LRLRGG ubiquitin peptide (left) and compound **5** (right)

To confirm that the screening hits bound at the C-terminal ubiquitin-binding pocket, and to guide future optimization efforts, we attempted soaking apo USP5 ZnF-UBD crystals with ligands but were unsuccessful. Co-crystallization attempts with the most potent compounds produced crystal structures with **1** (PDB: 6DXT), **5** (PDB: 6NFT) and **7** (PDB: 6DXH) (Figure 2). Compound **1** was not fully resolved by electron density, and modeled for only one of the two USP5 ZnF-UBD molecules in the crystallographic asymmetric unit. The carboxylate group of the ligands recapitulates interactions observed with the ubiquitin C-terminal di-glycine ^11^, and is engaged in three direct or water-mediated hydrogen bonds with surrounding residues. An aromatic ring π-stacks with Tyr259, but we did not observe a conformational rearrangement of Arg221 that could have further increased stacking interactions, as seen with the corresponding HDAC6 ZnF-UBD arginine residue, Arg1155. The absence of π-stacking interactions may be due to the absence of an extended polyaromatic system in the ligand. The side chain of Arg221 is engaged in a hydrogen-bond with the 1,3,4- oxadiazole ring of compound **1** and the carbonyl oxygens of compounds **5** and **7**. Interestingly, binding of compound **7** (PDB: 6DXH) induces the remodeling of a loop in the vicinity of the ligand (residues 221-230), which significantly opens up the binding pocket, and may offer a distinct vector for ligand optimization (Figure 3).

**Figure 2.**
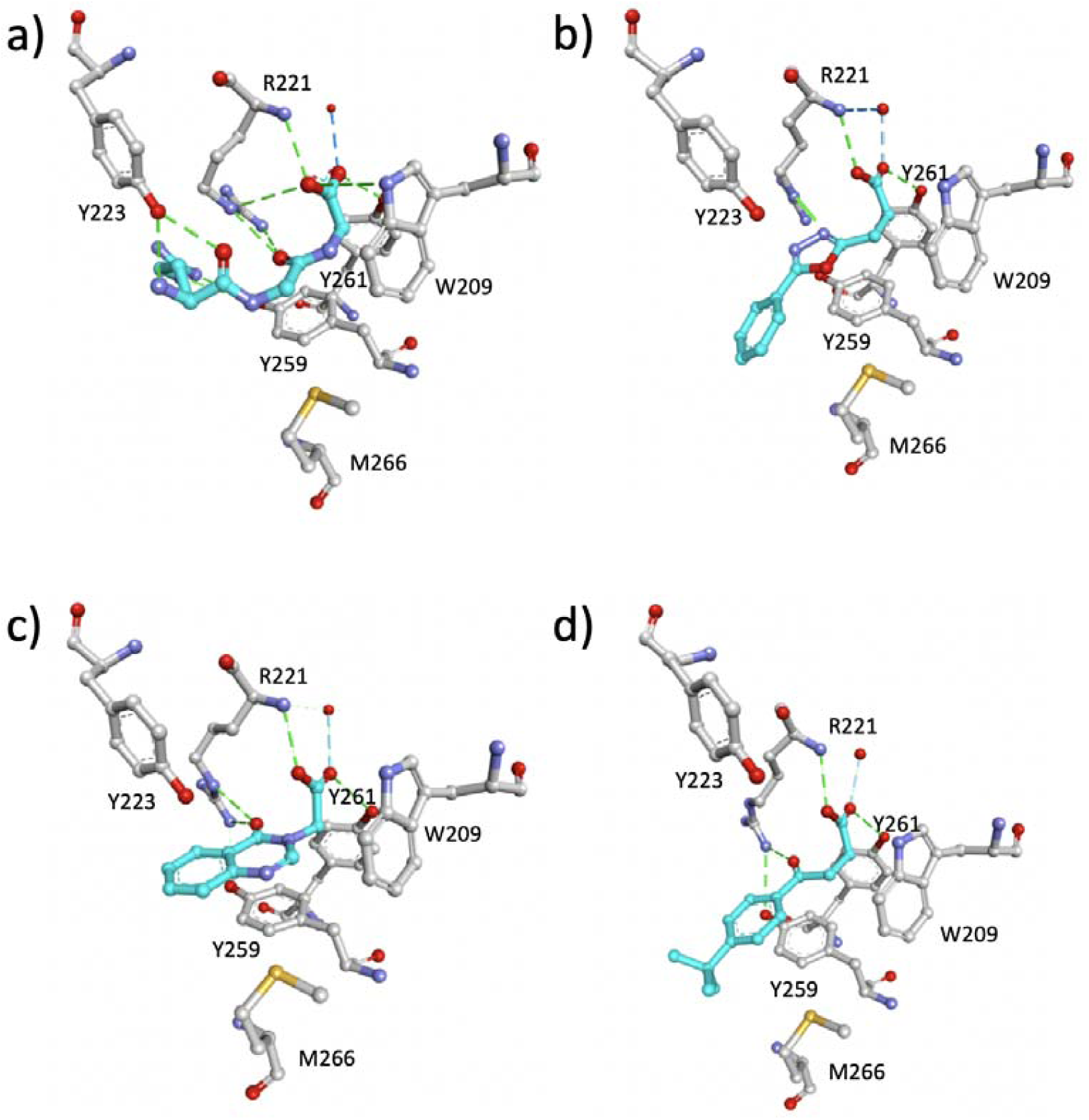
Crystal structure of USP5 ZnF-UBD in complex with a) C-terminal ubiquitin peptide (PDB: 2G45) b) **1** (PDB: 6DXT) c) **5** (PDB: 6NFT) d) **7** (PDB: 6DXH). Binding pocket residue (grey), ligands (cyan). Hydrogen bonds are shown as dotted green lines.

**Figure 3.**
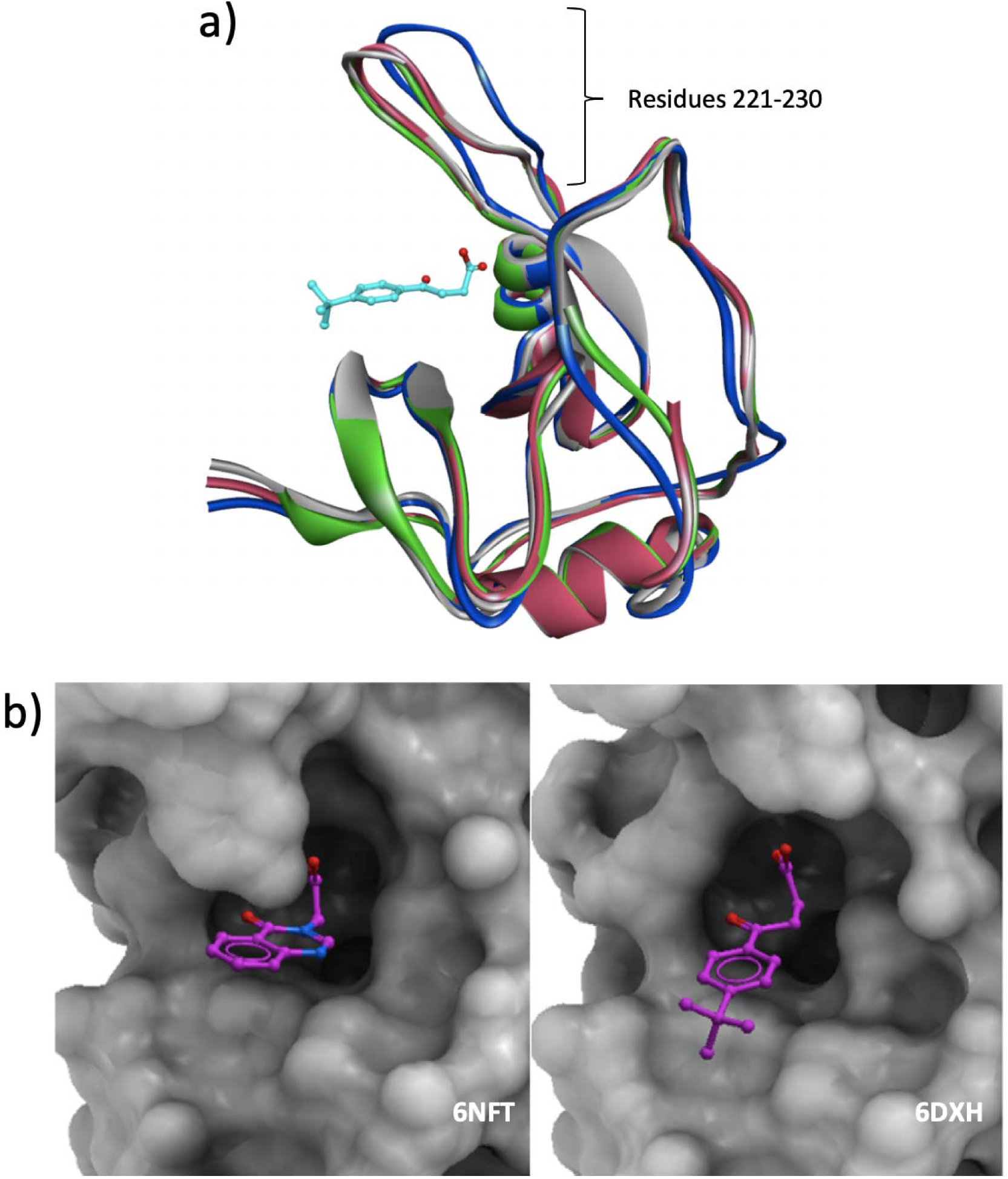
a) Superimposed crystal structure of USP5 ZnF-UBD (PDB: 2G43) (grey) in the apo state and bound to **1** (PDB: 6DXT) (magenta), **5** (PDB: 6NFT) (green), **7** (PDB:6DXH) (blue) highlighting the shifted loop in the compound **7** complex structure b) Surface representation of USP5 ZnF-UBD (grey) in complex with **5** and **7** (purple) (PDB: 6NFT and 6DXH)

### Hit Expansion

A substructure search of commercial libraries using the chemical scaffolds of compounds **1, 5**, and **7** found 399, 657 and 2450 analogues, respectively. The resulting libraries were docked to the USP5 ZnF-UBD crystallographic conformers corresponding to the respective ligand scaffold using Glide^38^. After visual inspection 30, 14 and 39 analogues of **1, 5**, and **7**, respectively, were selected and re-scored using free energy perturbation (FEP) (Desmond Molecular Dynamics System; D.E. Shaw Research: New York, NY, 2018). Compounds were clustered based on their chemical similarity, and physical samples were obtained for a selection based on high docking score and low molecular weight. 22 compounds were tested using an SPR assay (Table 2). Changes in the 1,3,4-oxadiazole dioxazole ring of **1**, which forms a critical hydrogen bond with the side chain of Arg221, were not tolerated (**12, 13** and **14**), and only analogs **18** and **19**, where a halogen atom decorates the phenyl ring showed a minor improvement in potency, with K_D_ values of 280 and 270 µM respectively. Among the 8 analogues of **5** (LE=0.33), two had improved potency; installing methoxy groups on the benzene ring (compound **22**) resulted in a K_D_ of 110 ± 10 µM, while adding a methyl group on the aliphatic carbon of the carboxylate tail (compound **21**) resulted in a K_D_ of 60 ± 8 µM and a good ligand efficiency of 0.36. Encouraged by this result, we subsequently ordered compounds **26, 26, 28** and **29**. Comparing **21** with **28** and **29** reveals that the pocket where the methyl group in **21** binds is very tight and does not tolerate larger hydrophobic groups. We solved the crystal structure of **21** in complex with USP5 ZnF-UBD, revealing that the added methyl group optimally occupies a hydrophobic cavity lined by Trp209, and a crystal packing similar to **7** with a shift in the loop orientation (Figure 4). Finally, one analogue of compound **7** (K_D_=170 ± 50 µM) had an improved binding affinity over the parent molecule. Compound **37** resulted in a K_D_ of 80 ± 1 µM, suggesting an i-propyl at the para position of the benzene ring is preferred over a t-butyl moiety. The preliminary structure activity relationship emerging from this work will serve as a framework for improvement of the current chemical series and exploration of novel ligand scaffolds. In particular, we believe that fragment **21** and **37** would be good starting points for future optimization efforts.

**Table 2.**
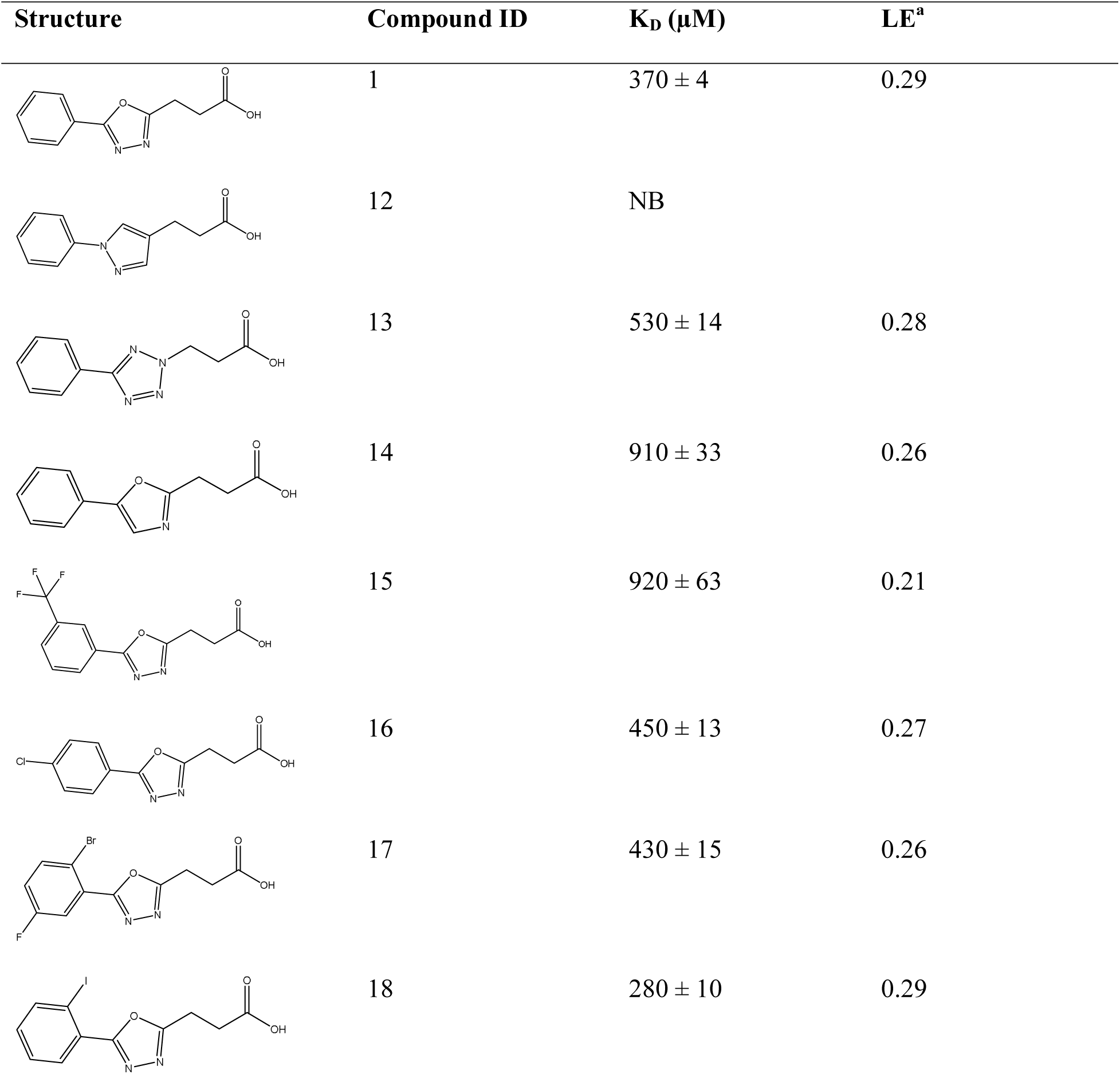
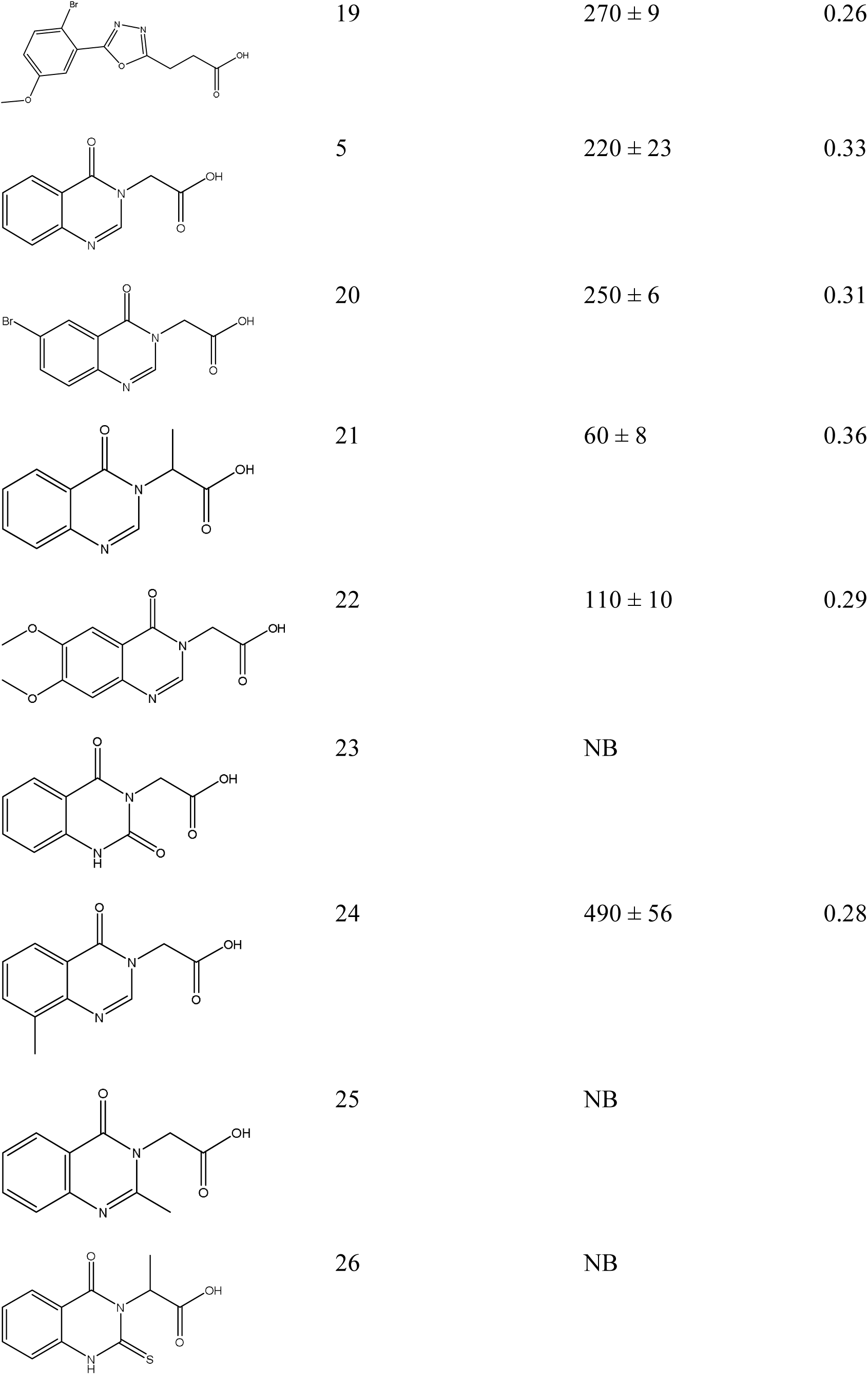
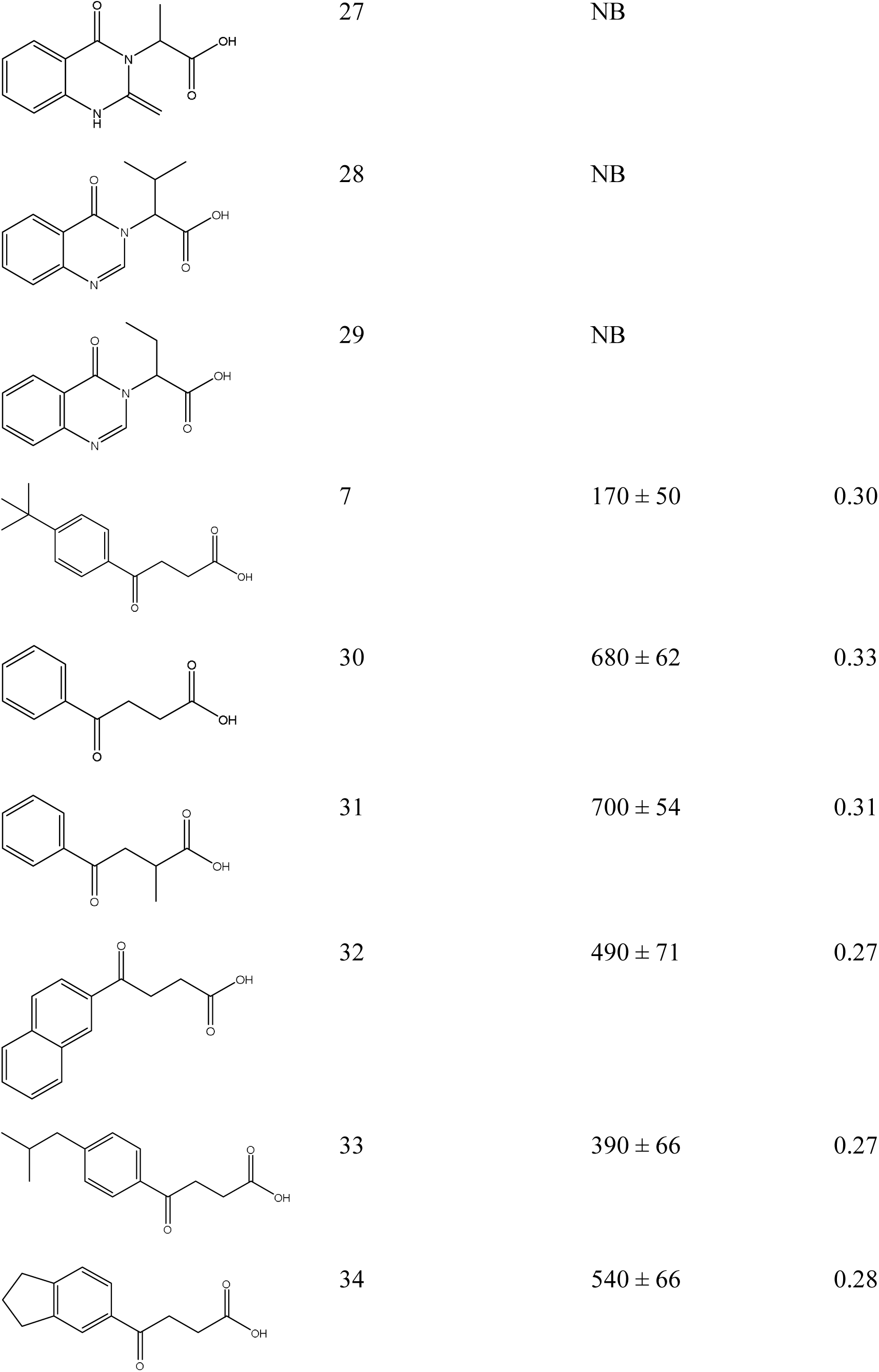
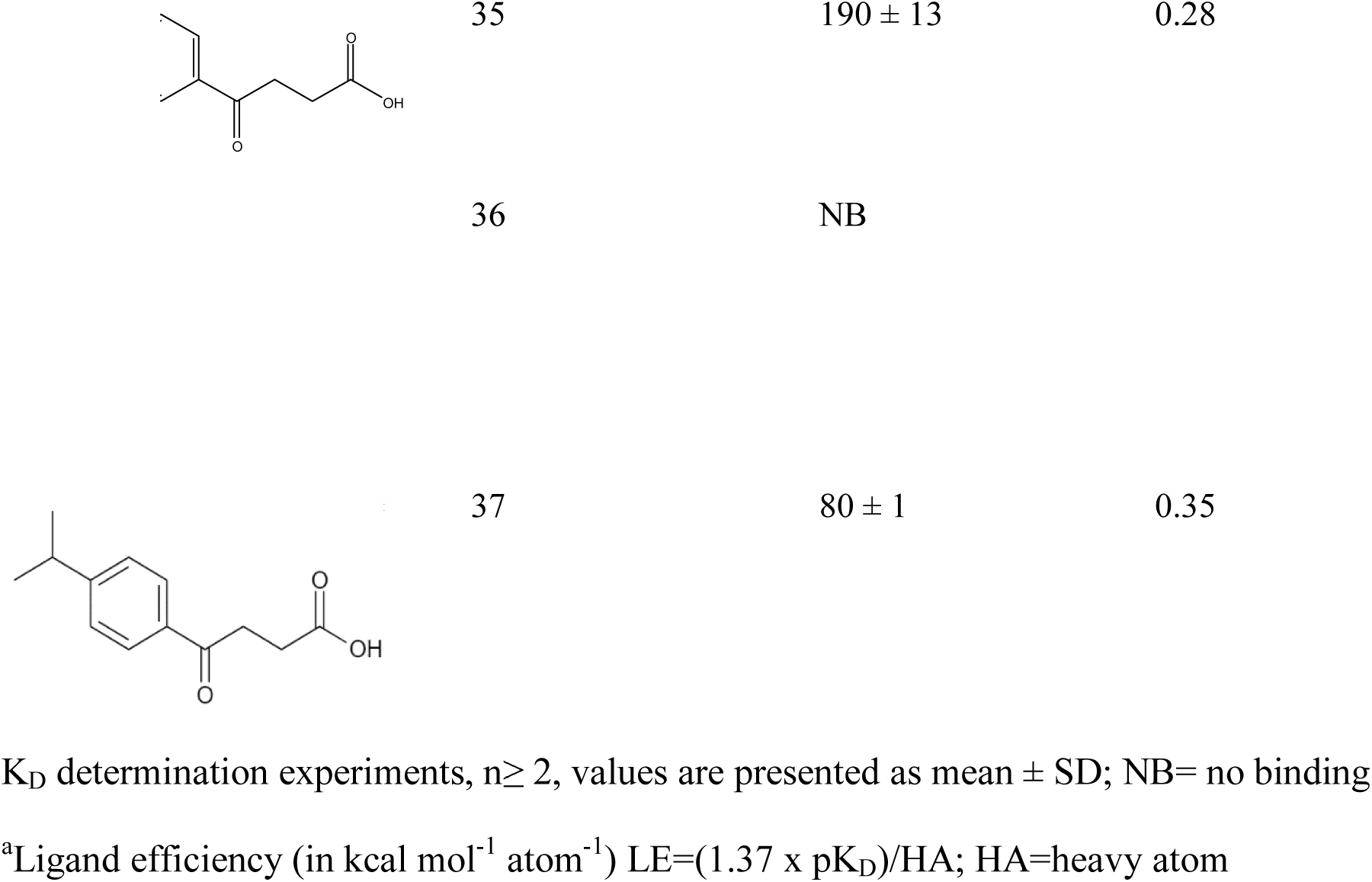
SPR K_D_ of Hit Expansion Compounds

**Figure 4.**
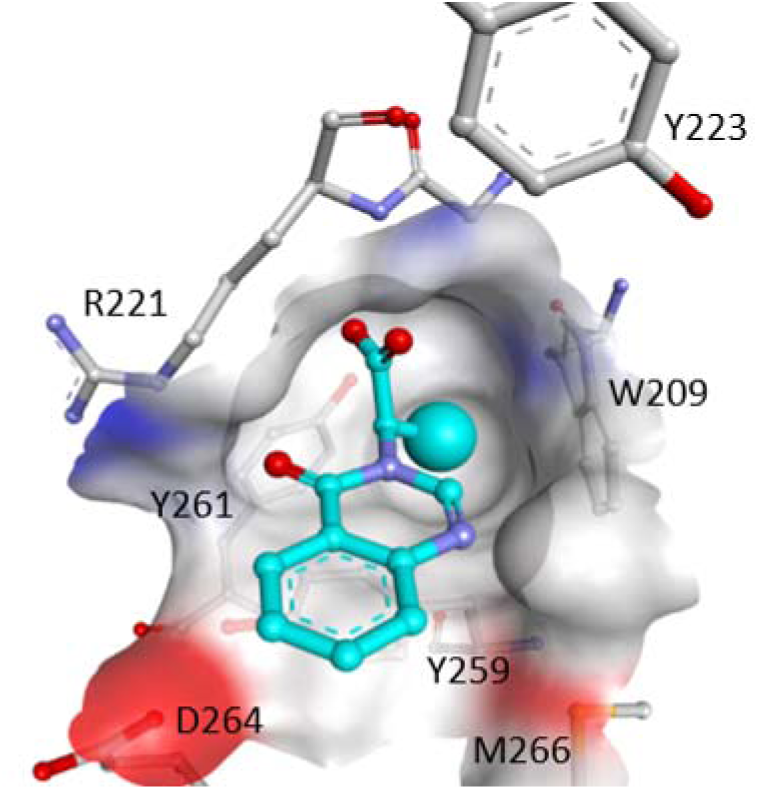
Co-crystal structure of USP5 ZnF-UBD (grey) in complex with **21** (cyan). A methyl group optimally exploiting a buried hydrophobic cavity is highlighted (PDB: 6P9G). The surface of the binding pocket is colored by binding properties: blue: hydrogen-bond donor, red: hydrogen-bond acceptor, grey: hydrophobic.

## Conclusion

Here, we report a combination of computational screens, binding assays, and crystal structures for the discovery of ligands targeting the ZnF-UBD of USP5. This work will serve as a platform to optimize the chemical series presented here, and potentially identify novel scaffolds, with the aim of developing potent inhibitors that are selective over ZnF-UBDs found in others USPs and HDAC6.

## Experimental Section

### Primary Virtual Screening

Ligprep (Schro□dinger Release 2018-2: LigPrep, Schro□dinger, LLC, New York, NY, 2018) and ICM ^37^ were used to prepare 3-dimensional models of 7680 compounds from commercial vendors and 1800 “in house” compounds. Protonation states at pH 7.4 ± 0.2 were generated with EpiK^39^ (Schro□dinger Release 2018-2: EpiK, Schro□dinger, LLC, New York, NY, 2018). The library compounds were docked using either ICM-Pro ^37^ or Glide SP^38^ (Schro□dinger Release 2018-2: Glide, Schro□dinger, LLC, New York, NY, 2018) to each of three representations of the USP5 ZnF-UBD: (1) a crystal conformer based on PDB entry 2G45 ^11^ including water molecule W404, which hydrogen-bonds to residues Arg221 and Asn207, referred to as “+W404”; (2) the same crystal conformer without that water molecule, “-W404”; (3) a simulated “stacked” receptor, obtained by re-modelling the side chain conformations for Arg221 and Tyr223 based on the stacked conformation of HDAC6 ZnF-UBD in complex with various inhibitors. After ICM docking with +W404, -W404 and “stacked” receptor representations, the 138, 89, and 139 best-scoring compounds were visually inspected, respectively; 28 compounds occurred in all three of the ICM best-scoring selections. After docking with the +404, −404 and “stacked” receptor representations, the 97, 103 and 162 best-scoring compounds were visually inspected, respectively. 36 compounds occurred in all three of the Glide best-scoring selections. 96 compounds were selected during visual inspection based on the subjective plausibility of the predicted binding pose and properties of compound chemical scaffolds. 33 compounds were ordered.

### Cloning, Protein Expression and Purification

DNA encoding USP5^171-290^ was sub-cloned into a modified pET28 vector encoding a TEV cleavable N-terminal His6-tag (pET28-MHL) and a modified vector with an N-terminal AviTag for biotinylation and C-terminal His6-tag (p28BIOH-LIC) using a ligation-independent InFusion cloning kit (ClonTech) and verified by DNA sequencing. Proteins were expressed in BL21 (DE3) Codon Plus RIL *Escherichia coli*. Cultures were grown in M9 minimal media, supplemented with 50 µM ZnSO_4_ for pET28-MHL and 10 µg/mL biotin for USP5^171-290^ p28BIOH-LIC expression. For USP5^171-290^ (pET28-MHL) with 5-fluoro-tryptophan (5FW) labels (Sigma Aldrich), cultures were grown in defined media ^40^. Expression cultures were induced at OD_600_ ∼0.6 with 0.5 mM IPTG overnight at 15□C. Proteins were purified using nickel-nitriloacetic acid (Ni-NTA) agarose resin, and the tag was removed by TEV for USP5^171-290^ pET28-MHL. Uncleaved proteins and TEV were removed by a second pass with Ni-NTA resin (Qiagen). Proteins were concentrated and further purified using gel filtration (S75 16/60, GE Healthcare) and ion-exchange (Hi-Load Q column, GE Healthcare) for the 5FW-labeled protein. The incorporation of the 5FW label was verified by mass spectrometry. The final concentration of purified proteins was 7-30 mg/mL as measured by UV absorbance at 280 nm.

### Crystallization

Crystallization solutions of 12 mg/mL of tag-free USP5^171-290^ (pET28-MHL) and 1% (v/v) and 4.5% (v/v) of a 200 mM DMSO-solubilized stock of compounds were prepared. Sparse-matrix crystallization experiments yielded crystals in different conditions for each USP5 ZnF-UBD-compound complex. 1:2.5 USP5^171-290^: compound **1** was co-crystallized with 1:1 protein/compound: mother liquor comprising 1.5 M ammonium sulfate, 0.1 M bis-tris pH 7.0. 1:2.5 USP5^171-290^: compound **5** was co-crystallized with 1:1 protein/compound: mother liquor comprising 2 M sodium/potassium phosphate pH 7.0. 1:5 USP5^171-290^: compound **7** was co-crystallized with 1:1 protein/compound: mother liquor comprising 1.5 M ammonium sulfate, 0.1 M bis-tris pH 7.0. 1:2.5 USP5^171-290^: compound **21** was co-crystallized with 1:1 protein/compound: mother liquor comprising 1.75 M ammonium sulfate, 0.2 M sodium acetate, 0.1 M sodium cacodylate pH 6.8. All co-crystals were cryo-protected with mother liquor supplemented with 25% ethylene glycol (v/v) prior to mounting and cryo-cooling.

### Data Collection, Structure Determination and Refinement

X-ray diffraction data for USP5^171-290^ co-crystals with **1, 5, 7, 21** were collected at 100 K at a Rigaku FR-E Superbright copper source. Diffraction data were processed with the XDS ^41^ and AIMLESS ^42^, using the Xia2 interfaces ^43^ for complex **1, 5** and **7**. Structures for compounds **1** and **5** were solved by direct refinement of protein chains A and B of the isomorphous PDB entry 2G43. The complex structure for compound **7** was solved by molecular replacement with Phaser ^44^ and coordinates from chain A of PDB entry 2G43. The co-crystal structure for compound **21** was solved by direct refinement of coordinates from PDB entry: 6DXH with the DIMPLE ^45^ script. Geomtry restraints for **21** were prepared with ACEDRG ^46^. Models were refined with cycles of COOT ^47^, for model building and visualization, REFMAC ^48^, for restrained refinement, and validated with MOLPROBITY ^49^.

### ^19^F NMR Spectroscopy

5FW USP5 USP5^171-290^ (at 25 µM) was buffered in 50 mM Tris pH 8, 150 mM NaCl, 1 mM TCEP, 0.005% Tween-20 (v/v), 5% D_2_O (v/v), 0.5% DMSO (v/v). For the binding screen, each compound was assayed individually (at 25:1 compound: protein). ^19^F spectra were acquired on a Bruker 600 MHz spectrometer equipped with a QCI cryoprobe. Each spectrum was acquired with 700-800 scans, an acquisition time of 90 ms, a recycle delay of 0.7 s and processed by applying an exponential window function (LB=10). The spectra was analyzed using software TopSpin (Bruker).

### Surface Plasmon Resonance

Studies were performed using a Biacore T200 (GE Health Sciences). Approximately 6000 response units (RU) of biotinylated USP5^171-290^ was coupled to flow cells of a SA chip per manufacturer’s protocol, and an empty flow cell used for reference subtraction. Two-fold serial dilutions were prepared in 20 mM Hepes pH 7.4, 150 mM NaCl, 0.005% Tween-20 (v/v), 1% DMSO (v/v). K_D_ determination experiments were performed using multi-cycle kinetics with 35 s contact time, and 30 µL/min flow rate at 20 □C. K_D_ values were calculated using steady state affinity fitting with the Biacore T200 evaluation software.

### Substructure search

Substructure search was run against ∼45 million commercial compounds from Molport (MolPort, version 2.92, 2019) using SMARTS strings generated in MarvinSketch (ChemAxon), with constraints on all aromatic rings and the carboxylate moiety. Similarity search against ∼6 million commercial compounds from Emolecules (Emolecules Inc, 2019), was run using SMILES with a similarity index of 0.8.

### Virtual hit expansion

LigPrep (Schro□dinger Release 2018-2: LigPrep, Schro□dinger, LLC, New York, NY, 2018) was used to prepare the ligands using default settings. The library designed by the substructure search was docked to the USP5 ZnF-UBD structure for each respective complex (**1, 5, 7**) using Glide^38^ (Schro□dinger Release 2018-2: Glide, Schro□dinger, LLC, New York, NY, 2018). Three hydrogen-bond constraints between the carboxylate moiety of the ligand and NH of Arg221 backbone, NH of Arg221 side chain and OH of Tyr261. Docked compounds were clustered in ICM ^37^ and selected based on docking scores, complexity and visual inspection of docked poses.

### Free Energy Perturbation

All FEP calculations were conducted with the academic LigandFEP methodology of Desmond (Desmond Molecular Dynamics System; D. E. Shaw Research: New York, NY, 2017; ^50^ using the OPLS_2005 force field ^51^. The systems were solvated in an orthogonal box of SPC water molecules with buffer width of 5 Å for the complex and 10 Å for the solvent simulations. The full systems were relaxed and equilibrated using the default Desmond protocol, consisting of: (i) a minimization using a Brownian dynamics NVT integrator for 100ps with the solute molecules restrained (50 kcal/mol/Å2), (ii) 12 ps simulation in the NVT ensemble, keeping the restraints and temperature at 10 K, (iii) 12 ps simulation in the NPT ensemble, keeping restraints and temperature at 10 K, (iv) 24 ps simulation in the NPT simulation with solute heavy atom restraints at 300 K, and (v) 240 ps simulation in the NPT ensemble at 300 K without restraints. Production simulations in the NPT ensemble lasted 5 ns for both the complex and the solvent systems. A total of 12 λ windows were used for all calculations. The free energy differences between the initial and final states were calculated using the Bennett acceptance ratio (BAR) method ^52^, and the errors estimated using bootstrapping ^53^.

### Liquid chromatography-mass spectrometry (LCMS)

Chromatographic analyses were carried out on an ACQUITY UPLC BEH C18 (2.1 × 50 mm, 1.7 µm) column. The mobile phase was 0.1% formic acid in water (solvent A) and 0.1% formic acid in acetonitrile (solvent B) at a flow rate of 0.4 mL/min. A gradient starting at 95% solvent A going to 5% in 4.5 min., holding for 0.5 min., going back to 95% in 0.5 min and equilibrating the column for 1 min was employed. A Waters Xevo QTof or a Waters SYNAPT G2-S MS equipped with an atmospheric pressure ionization source was used for MS analysis. MassLynx 4.1 was used for data analysis.

### Accession Codes

**1** (6DXT), **5** (6NFT), **7** (6DXH), **21** (6P9G)

Authors will release atomic coordinates and experimental data upon article publication.

## Author Information

### Author Contributions

M.M. and M.S. wrote the manuscript. I.F. and R.F.F. completed preliminary docking studies. R.H., and M.M. performed the crystallographic calculations and model refinement for X-ray crystal structures with **1, 5**, and **7**. W.T. performed the crystallographic calculations and model refinement of X-ray structure with compound **21**. M.M. and S.H tested compounds. M.M. completed hit expansion. C.H.A, R.H. and M.S. advised throughout the project.

### Notes

The authors declare no competing financial interest.

## Acknowledgments

The SGC is a registered charity (number 1097737) that receives funds from AbbVie, Bayer Pharma AG, Boehringer Ingelheim, Canada Foundation for Innovation, Eshelman Institute for Innovation, Genome Canada through Ontario Genomics Institute [OGI-055], Innovative Medicines Initiative (EU/EFPIA) [ULTRA-DD grant no. 115766], Janssen, Merck KGaA, Darmstadt, Germany, MSD, Novartis Pharma AG, Innovation and Science (MRIS), Pfizer, São Paulo Research Foundation-FAPESP, Takeda, and Wellcome. M.S. gratefully acknowledges support from NSERC [Grant RGPIN-2019-04416]. C.H.A. gratefully acknowledges support from NSERC [RGPIN-2015-05939]. R.H. is the recipient of the Huntington’s Disease Society of America Berman Topper Career Development Fellowship. We are grateful to D.E. Shaw Research for providing us with an academic license for their Desmond. We thank Peter Brown and OICR for analytical LCMS of compounds.

## Abbreviations Used

BAR: Bennett acceptance ratio;
DUB: deubiquitinase;
FEP: free energy perturbation;
K_D_: dissociation constant;
SAR: structure activity relationship;
SD: standard deviation;
SPR: surface plasmon resonance;
Ub: ubiquitin;
UBA: ubiquitin associated domain;
UBP: ubiquitin-binding domain;
USP: ubiquitin specific protease;
ZnF-UBD: zinc finger ubiquitin-binding domain;
5FW: 5-fluoro-tryptophan

## Supporting Information

- USP5 co-crystal structure 6DXT (CIF)
- USP5 co-crystal structure 6DXT-sf (CIF)
- USP5 co-crystal structure 6NFT (CIF)
- USP5 co-crystal structure 6NFT-sf (CIF)
- USP5 co-crystal structure 6DXH (CIF)
- USP5 co-crystal structure 6DXH-sf (CIF)
- USP5 co-crystal structure 6P9G (CIF)
- USP5 co-crystal structure 6P9G-sf (CIF)
- X-ray crystallography data collection and refinement statistics and NMR screening data (PDF)
- Molecular formula strings (CSV)

## Table of Contents Graphic

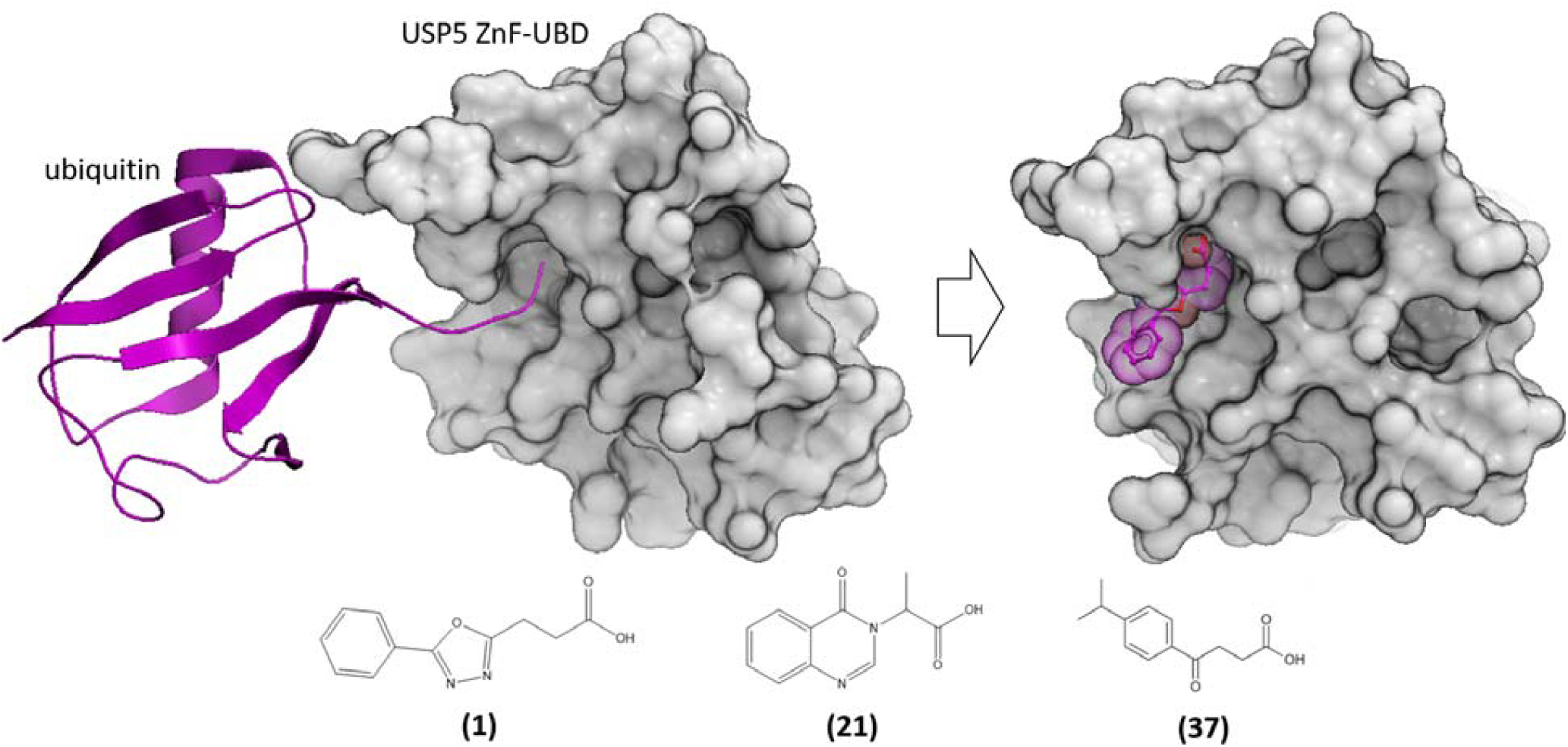

## References

(1) Hershko and Ciechanover. The Ubiquitin System. Trends Biochem. Sci. 1998. https://doi.org/10.1016/S0968-0004(97)01122-5.

(2) Yuan, T.; Yan, F.; Ying, M.; Cao, J.; He, Q.; Zhu, H.; Yang, B. Inhibition of Ubiquitin-Specific Proteases as a Novel Anticancer Therapeutic Strategy. Front. Pharmacol. 2018. https://doi.org/10.3389/fphar.2018.01080.

(3) Reyes-Turcu, F. E.; Ventii, K. H.; Wilkinson, K. D. Regulation and Cellular Roles of Ubiquitin-Specific Deubiquitinating Enzymes. Annu. Rev. Biochem. 2009. https://doi.org/10.1146/annurev.biochem.78.082307.091526.

(4) Clague, M. J.; Heride, C.; Urbé, S. The Demographics of the Ubiquitin System. Trends in Cell Biology. 2015. https://doi.org/10.1016/j.tcb.2015.03.002.

(5) Swatek, K. N.; Komander, D. Ubiquitin Modifications. Cell Research. 2016. https://doi.org/10.1038/cr.2016.39.

(6) Yau, R.; Rape, M. The Increasing Complexity of the Ubiquitin Code. Nature Cell Biology. 2016. https://doi.org/10.1038/ncb3358.

(7) Walczak, H.; Iwai, K.; Dikic, I. Generation and Physiological Roles of Linear Ubiquitin Chains. BMC Biology. 2012. https://doi.org/10.1186/1741-7007-10-23.

(8) Wilkinson, K. D.; Tashayev, V. L.; O’Connor, L. B.; Larsen, C. N.; Kasperek, E.; Pickart, C. M. Metabolism of the Polyubiquitin Degradation Signal: Structure, Mechanism, and Role of Isopeptidase T. Biochemistry 1995. https://doi.org/10.1021/bi00044a032.

(9) Welchman, R. L.; Gordon, C.; Mayer, R. J. Ubiquitin and Ubiquitin-like Proteins as Multifunctional Signals. Nature Reviews Molecular Cell Biology. 2005. https://doi.org/10.1038/nrm1700.

(10) Clague, M. J.; Urbé, S.; Komander, D. Breaking the Chains: Deubiquitylating Enzyme Specificity Begets Function. Nat. Rev. Mol. Cell Biol. 2019. https://doi.org/10.1038/s41580-019-0099-1.

(11) Reyes-Turcu, F. E.; Horton, J. R.; Mullally, J. E.; Heroux, A.; Cheng, X.; Wilkinson, K. D. The Ubiquitin Binding Domain ZnF UBP Recognizes the C-Terminal Diglycine Motif of Unanchored Ubiquitin. Cell 2006. https://doi.org/10.1016/j.cell.2006.02.038.

(12) Avvakumov, G. V.; Walker, J. R.; Xue, S.; Allali-Hassani, A.; Asinas, A.; Nair, U. B.; Fang, X.; Zuo, X.; Wang, Y. X.; Wilkinson, K. D.; et al. Two ZnF-UBP Domains in Isopeptidase T (USP5). Biochemistry 2012. https://doi.org/10.1021/bi200854q.

(13) Amerik, A. Y.; Hochstrasser, M. Mechanism and Function of Deubiquitinating Enzymes. Biochimica et Biophysica Acta - Molecular Cell Research. 2004. https://doi.org/10.1016/j.bbamcr.2004.10.003.

(14) Dang, L. C.; Melandri, F. D.; Stein, R. L. Kinetic and Mechanistic Studies on the Hydrolysis of Ubiquitin C-Terminal 7-Amido-4-Methylcoumarin by Deubiquitinating Enzymes. Biochemistry 1998. https://doi.org/10.1021/bi9723360.

(15) Dayal, S.; Sparks, A.; Jacob, J.; Allende-Vega, N.; Lane, D. P.; Saville, M. K. Suppression of the Deubiquitinating Enzyme USP5 Causes the Accumulation of Unanchored Polyubiquitin and the Activation of P53. J. Biol. Chem. 2009. https://doi.org/10.1074/jbc.M805871200.

(16) Liu, Y.; Wang, W. M.; Zou, L. Y.; Li, L.; Feng, L.; Pan, M. Z.; Lv, M. Y.; Cao, Y.; Wang, H.; Kung, H. F.; et al. Ubiquitin Specific Peptidase 5 Mediates Histidine-Rich Protein Hpn Induced Cell Apoptosis in Hepatocellular Carcinoma through P14-P53 Signaling. Proteomics 2017. https://doi.org/10.1002/pmic.201600350.

(17) Kaistha, B. P.; Krattenmacher, A.; Fredebohm, J.; Schmidt, H.; Behrens, D.; Widder, M.; Hackert, T.; Strobel, O.; Hoheisel, J. D.; Gress, T. M.; et al. The Deubiquitinating Enzyme USP5 Promotes Pancreatic Cancer via Modulating Cell Cycle Regulators. Oncotarget 2017.

(18) Wang, S.; Juan, J.; Zhang, Z.; Du, Y.; Xu, Y.; Tong, J.; Cao, B.; Moran, M. F.; Zeng, Y.; Mao, X. Inhibition of the Deubiquitinase USP5 Leads to C-Maf Protein Degradation and Myeloma Cell Apoptosis. Cell Death Dis. 2017. https://doi.org/10.1038/cddis.2017.450.

(19) Ma, X.; Qi, W.; Pan, H.; Yang, F.; Deng, J. Overexpression of USP5 Contributes to Tumorigenesis in Non-Small Cell Lung Cancer via the Stabilization of β-Catenin Protein. Am. J. Cancer Res. 2018.

(20) GarcíAllali-Caballero, A.; Gadotti, V. M.; Stemkowski, P.; Weiss, N.; Souza, I. A.; Hodgkinson, V.; Bladen, C.; Chen, L.; Hamid, J.; Pizzoccaro, A.; et al. The Deubiquitinating Enzyme USP5 Modulates Neuropathic and Inflammatory Pain by Enhancing Cav3.2 Channel Activity. Neuron 2014. https://doi.org/10.1016/j.neuron.2014.07.036.

(21) Allali-Caballero, A.; Gadotti, V. M.; Chen, L.; Zamponi, G. W. A Cell-Permeant Peptide Corresponding to the CUBP Domain of USP5 Reverses Inflammatory and Neuropathic Pain. Mol. Pain 2016. https://doi.org/10.1177/1744806916642444.

(22) Stemkowski, P. L.; Garcia-Caballero, A.; Gadotti, V. M.; M’Dahoma, S.; Chen, L.; Souza, A.; Zamponi, G. W. Identification of Interleukin-1 Beta as a Key Mediator in the Upregulation of Cav3.2–USP5 Interactions in the Pain Pathway. Mol. Pain 2017. https://doi.org/10.1177/1744806917724698.

(23) Liu, Q.; Wu, Y.; Qin, Y.; Hu, J.; Xie, W.; Xiao-Feng Qin, F.; Cui, J. Broad and Diverse Mechanisms Used by Deubiquitinase Family Members in Regulating the Type i Interferon Signaling Pathway during Antiviral Responses. Sci. Adv. 2018. https://doi.org/10.1126/sciadv.aar2824.

(24) Ovaa, H.; Kessler, B. M.; Rolen, U.; Galardy, P. J.; Ploegh, H. L.; Masucci, M. G. Activity-Based Ubiquitin-Specific Protease (USP) Profiling of Virus-Infected and Malignant Human Cells. Proc. Natl. Acad. Sci. 2004. https://doi.org/10.1073/pnas.0308411100.

(25) Nakajima, S.; Lan, L.; Wei, L.; Hsieh, C. L.; Rapić-Otrin, V.; Yasui, A.; Levine, A. S. Ubiquitin-Specific Protease 5 Is Required for the Efficient Repair of DNA Double-Strand Breaks. PLoS One 2014. https://doi.org/10.1371/journal.pone.0084899.

(26) Xie, X.; Matsumoto, S.; Endo, A.; Fukushima, T.; Kawahara, H.; Saeki, Y.; Komada, M. Deubiquitylases USP5 and USP13 Are Recruited to and Regulate Heat-Induced Stress Granules through Their Deubiquitylating Activities. J. Cell Sci. 2018. https://doi.org/10.1242/jcs.210856.

(27) Meyers, R. M.; Bryan, J. G.; McFarland, J. M.; Weir, B. A.; Sizemore, A. E.; Xu, H.; Dharia, N. V.; Montgomery, P. G.; Cowley, G. S.; Pantel, S.; et al. Computational Correction of Copy Number Effect Improves Specificity of CRISPR-Cas9 Essentiality Screens in Cancer Cells. Nat. Genet. 2017. https://doi.org/10.1038/ng.3984.

(28) Behan, F. M.; Iorio, F.; Picco, G.; Gonçalves, E.; Beaver, C. M.; Migliardi, G.; Santos, R.; Rao, Y.; Sassi, F.; Pinnelli, M.; et al. Prioritization of Cancer Therapeutic Targets Using CRISPR–Cas9 Screens. Nature 2019. https://doi.org/10.1038/s41586-019-1103-9.

(29) Depmap Broad. DepMap Achilles 19Q1 Public. doi:10.6084/m9.figshare.7655150.

(30) Gavory, G.; O’Dowd, C. R.; Helm, M. D.; Flasz, J.; Arkoudis, E.; Dossang, A.; Hughes, C.; Cassidy, E.; McClelland, K.; Odrzywol, E.; et al. Discovery and Characterization of Highly Potent and Selective Allosteric USP7 Inhibitors. Nat. Chem. Biol. 2018. https://doi.org/10.1038/nchembio.2528.

(31) Kategaya, L.; Di Lello, P.; Rougé, L.; Pastor, R.; Clark, K. R.; Drummond, J.; Kleinheinz, T.; Lin, E.; Upton, J. P.; Prakash, S.; et al. USP7 Small-Molecule Inhibitors Interfere with Ubiquitin Binding. Nature 2017. https://doi.org/10.1038/nature24006.

(32) Liang, Q.; Dexheimer, T. S.; Zhang, P.; Rosenthal, A. S.; Villamil, M. A.; You, C.; Zhang, Q.; Chen, J.; Ott, C. A.; Sun, H.; et al. A Selective USP1-UAF1 Inhibitor Links Deubiquitination to DNA Damage Responses. Nat. Chem. Biol. 2014. https://doi.org/10.1038/nchembio.1455.

(33) Turnbull, A. P.; Ioannidis, S.; Krajewski, W. W.; Pinto-Fernandez, A.; Heride, C.; Martin, A. C. L.; Tonkin, L. M.; Townsend, E. C.; Buker, S. M.; Lancia, D. R.; et al. Molecular Basis of USP7 Inhibition by Selective Small-Molecule Inhibitors. Nature 2017. https://doi.org/10.1038/nature24451.

(34) Wang, Y.; Jiang, Y.; Ding, S.; Li, J.; Song, N.; Ren, Y.; Hong, D.; Wu, C.; Li, B.; Wang, F.; et al. Small Molecule Inhibitors Reveal Allosteric Regulation of USP14 via Steric Blockade. Cell Res. 2018. https://doi.org/10.1038/s41422-018-0091-x.

(35) Ferreira De Freitas, R.; Harding, R. J.; Franzoni, I.; Ravichandran, M.; Mann, M. K.; Ouyang, H.; Lautens, M.; Santhakumar, V.; Arrowsmith, C. H.; Schapira, M. Identification and Structure-Activity Relationship of HDAC6 Zinc-Finger Ubiquitin Binding Domain Inhibitors. J. Med. Chem. 2018. https://doi.org/10.1021/acs.jmedchem.8b00258.

(36) Harding, R. J.; Ferreira De Freitas, R.; Collins, P.; Franzoni, I.; Ravichandran, M.; Ouyang, H.; Juarez-Ornelas, K. A.; Lautens, M.; Schapira, M.; Von Delft, F.; et al. Small Molecule Antagonists of the Interaction between the Histone Deacetylase 6 Zinc-Finger Domain and Ubiquitin. J. Med. Chem. 2017. https://doi.org/10.1021/acs.jmedchem.7b00933.

(37) Abagyan, R.; Totrov, M.; Kuznetsov, D. ICM—A New Method for Protein Modeling and Design: Applications to Docking and Structure Prediction from the Distorted Native Conformation. J. Comput. Chem. 1994. https://doi.org/10.1002/jcc.540150503.

(38) Halgren, T. A.; Murphy, R. B.; Friesner, R. A.; Beard, H. S.; Frye, L. L.; Pollard, W. T.; Banks, J. L. Glide: A New Approach for Rapid, Accurate Docking and Scoring. 2. Enrichment Factors in Database Screening. J. Med. Chem. 2004. https://doi.org/10.1021/jm030644s.

(39) Shelley, J. C.; Cholleti, A.; Frye, L. L.; Greenwood, J. R.; Timlin, M. R.; Uchimaya, M. Epik: A Software Program for PKa Prediction and Protonation State Generation for Drug-like Molecules. J. Comput. Aided. Mol. Des. 2007. https://doi.org/10.1007/s10822-007-9133-z.

(40) Gee, C. T.; Arntson, K. E.; Urick, A. K.; Mishra, N. K.; Hawk, L. M. L.; Wisniewski, A. J.; Pomerantz, W. C. K. Protein-Observed 19F-NMR for Fragment Screening, Affinity Quantification and Druggability Assessment. Nat. Protoc. 2016. https://doi.org/10.1038/nprot.2016.079.

(41) Kabsch, W. XDS. Acta Crystallogr. Sect. D Biol. Crystallogr. 2010. https://doi.org/10.1107/S0907444909047337.

(42) Evans, P. R.; Murshudov, G. N. How Good Are My Data and What Is the Resolution? Acta Crystallogr. Sect. D Biol. Crystallogr. 2013. https://doi.org/10.1107/s0907444913000061.

(43) Winter, G. Xia2: An Expert System for Macromolecular Crystallography Data Reduction. J. Appl. Crystallogr. 2010. https://doi.org/10.1107/S0021889809045701.

(44) McCoy, A. J.; Grosse-Kunstleve, R. W.; Adams, P. D.; Winn, M. D.; Storoni, L. C.; Read, R. J. Phaser Crystallographic Software. J. Appl. Crystallogr. 2007. https://doi.org/10.1107/s0021889807021206.

(45) Wojdyr, M.; Keegan, R.; Winter, G.; Ashton, A. DIMPLE - a Pipeline for the Rapid Generation of Difference Maps from Protein Crystals with Putatively Bound Ligands. Acta Crystallogr. Sect. A Found. Crystallogr. 2013. https://doi.org/10.1107/s0108767313097419.

(46) Long, F.; Nicholls, R. A.; Emsley, P.; Gražulis, S.; Merkys, A.; Vaitkus, A.; Murshudov, G. N. AceDRG□: A Stereochemical Description Generator for Ligands. Acta Crystallogr. Sect. D Struct. Biol. 2017. https://doi.org/10.1107/s2059798317000067.

(47) Emsley, P.; Lohkamp, B.; Scott, W. G.; Cowtan, K. Features and Development of \textit{Coot}. Acta Crystallogr D Biol Crystallogr 2010. https://doi.org/10.1107/S0907444910007493.

(48) Murshudov, G. N.; Skubák, P.; Lebedev, A. A.; Pannu, N. S.; Steiner, R. A.; Nicholls, R. A.; Winn, M. D.; Long, F.; Vagin, A. A. REFMAC5 for the Refinement of Macromolecular Crystal Structures. Acta Crystallogr. D. Biol. Crystallogr. 2011. https://doi.org/10.1107/S0907444911001314.

(49) Chen, V. B.; Arendall, W. B.; Headd, J. J.; Keedy, D. A.; Immormino, R. M.; Kapral, G. J.; Murray, L. W.; Richardson, J. S.; Richardson, D. C. MolProbity: All-Atom Structure Validation for Macromolecular Crystallography. Acta Crystallogr. Sect. D Biol. Crystallogr. 2010. https://doi.org/10.1107/S0907444909042073.

(50) Bowers, K. J.; Sacerdoti, F. D.; Salmon, J. K.; Shan, Y.; Shaw, D. E.; Chow, E.; Xu, H.; Dror, R. O.; Eastwood, M. P.; Gregersen, B. A.; et al. Molecular Dynamics---Scalable Algorithms for Molecular Dynamics Simulations on Commodity Clusters. In Proceedings of the 2006 ACM/IEEE conference on Supercomputing - SC ‘06; 2006. https://doi.org/10.1145/1188455.1188544.

(51) Banks, J. L.; Beard, H. S.; Cao, Y.; Cho, A. E.; Damm, W.; Farid, R.; Felts, A. K.; Halgren, T. A.; Mainz, D. T.; Maple, J. R.; et al. Integrated Modeling Program, Applied Chemical Theory (IMPACT). Journal of Computational Chemistry. 2005. https://doi.org/10.1002/jcc.20292.

(52) Bennett, C. H. Efficient Estimation of Free Energy Differences from Monte Carlo Data. J. Comput. Phys. 1976. https://doi.org/10.1016/0021-9991(76)90078-4.

(53) Paliwal, H.; Shirts, M. R. A Benchmark Test Set for Alchemical Free Energy Transformations and Its Use to Quantify Error in Common Free Energy Methods. J. Chem. Theory Comput. 2011. https://doi.org/10.1021/ct2003995.

