## Supporting Information for "Discovery of small molecule antagonists of the USP5 zinc finger ubiquitin-binding domain"

[*]Corresponding authors

**Contents**

Supplementary tables and figures p S3-S5

**Table S1**. Data collection and refinement statistics for USP5 co-crystal structures

| **PDB code** | **6DXT** | **6NFT** | **6DXH** | **6P9G** |
|---|---|---|---|---|
| Compound | 1 | 5 | 7 | 21 |
| Space group | C2 | C2 | I222 | I222 |
| a,b,c [Å] | 61.09,85.44,59.74 | 61.87,85.04,59.84 | 47.69,81.50,99.66 | 47.85,82.41,99.82 |
| α,β,γ [°] | 90.00,100.29,90.00 | 90.00,99.00,90.00 | 90.00,90.00,90.00 | 90.00,90.00,90.00 |
| Resolution limits [Å] | 30.05-1.95(2.00-1.95) | 26.94-1.65(1.68-1.65) | 31.54-2.00(2.05-2.00) | 43.15-1.84(1.88-1.84) |
| Rmerge | 0.038(0.218) | 0.084(0.916) | 0.054(0.451) | 0.068(1.791) |
| I/sigma | 16.2(4.8) | 10.7(1.6) | 23.0(4.4) | 19.7(1.5) |
| Completeness [%] | 95.2(92.9) | 99.2(97.7) | 99.4(98.3) | 99.9(99.2) |
| Multiplicity | 3.8(3.9) | 4.0(3.9) | 7.0(7.0) | 6.3(6.3) |
| Resolution [A] | 30.07-1.95 | 26.94-1.65 | 31.56-2.00 | 41.00-2.10 |
| No. Reflections used/free | 19928/1010 | 34601/1821 | 12742/679 | 11293/586 |
| Rwork/Rfree | 0.194/0.242 | 0.180/0.206 | 0.214/0.251 | 0.224/0.248 |
| No. atoms/B-factors [Å$^2$] | 1945/26.9 | 2136/19.3 | 981/36.0 | 936/43.0 |
| Protein | 1747/26.1 | 1830/18.1 | 902/35.7 | 893/43.3 |
| Inhibitor | 16/38.8 | 30/15.9 | 17/36.1 | 16/38.8 |
| Water | 170/33.6 | 245/28.4 | 54/41.2 | 13/34.5 |
| Others | 12/26.0 | 31/24.2 | 8/30.1 | 14/32.5 |
| rmsd bonds [Å]/ angles [°] | 0.011/1.5 | 0.011/1.7 | 0.012/1.6 | 0.010/1.7 |
| Molprobity Ramachandran favored/outliers [%] | 98.65/0.00 | 98.25/0.00 | 98.26/0.00 | 97.39/0.00 |

This table was prepared with PDB_EXTRACT, IOTBX and PHENIX software.

**Table S2**. Preliminary $^{19}$F NMR Screen and SPR data

| **SGC ID** | **Compound Structure** | **Manuscript ID** | **Peak 1 δ (ppm)** | **Δδ (ppm) [rel to control peak 1]** | **SPR $K_D$** |
|---|---|---|---|---|---|
| Control* | | | 118.20 | 0.00 | |
| DAT00000180a | 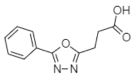 | 1 | 119.33 | 1.13 | 370 ± 4 |
| DAT00000183a | 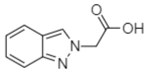 | 2 | 118.99 | 0.79 | 930 ± 78 |
| DAT00000187a | 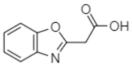 | 3 | 118.83 | 0.63 | 770 ± 49 |
| DAT00000190a | 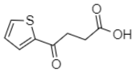 | 4 | 119.67 | 1.47 | 840 ± 26 |
| DAT00000194a | 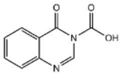 | 5 | 119.42 | 1.22 | 220 ± 23 |
| DAT00000198a | 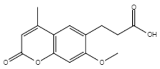 | 6 | 119.41 | 1.21 | 250 ± 6 |
| DAT00000201a | 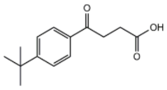 | 7 | 119.74 | 1.54 | 170 ± 50 |
| DAT00000202a | 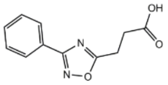 | 8 | 118.67 | 0.47 | NB |
| DAT00000203a | 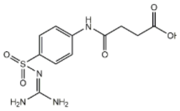 | 9 | 118.84 | 0.64 | NB |
| DAT00000208a | 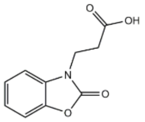 | 10 | 118.80 | 0.60 | NB |
| DAT00000212a | 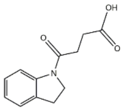 | 11 | 119.42 | 1.22 | NB |

*Control= USP5 ZnF-UBD only
NB=no binding

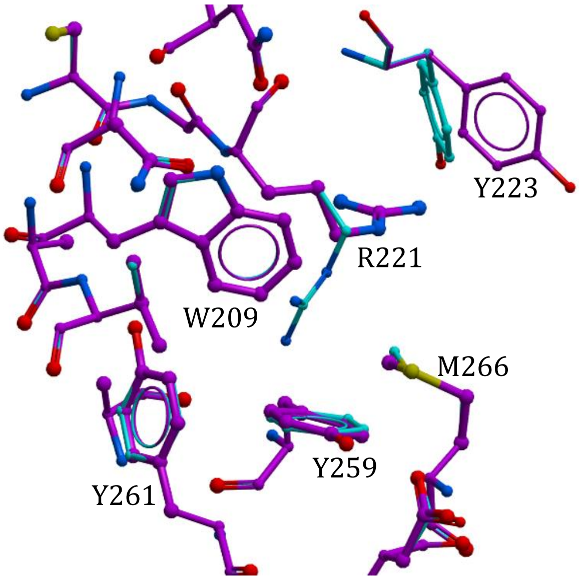


**Figure S1.** Superimposed crystal structure (PDB: 2G45) of USP5 ZnF-UBD (cyan) and modelled stacked conformation (purple) highlighting the different positioning of Arg221 and Tyr223.
